# Microbial Immunomodulatory Function is Altered in Inflammatory Bowel Disease Independent of Microbial Composition

**DOI:** 10.64898/2026.07.22.740186

**Authors:** Rabina Giri, Jing Jie Teh, Fan Zhang, Hayley Reed, Georgina L Hold, Paraic O Cuiv, Mark Morrison, Jakob Begun

## Abstract

Inflammatory bowel diseases (IBD) are chronic and relapsing immune-mediated conditions in which the NF-κB and STAT3 signalling pathways play central roles in pathogenesis. While gut microbiome dysbiosis is well described in IBD, its functional consequences, particularly microbial immunomodulatory function (MIF) remain poorly defined. Here, we integrated functional assays with metagenomic profiling to characterise microbiome-driven immune modulation in IBD.

Faecal water from non-IBD controls and IBD patients was assessed using NF-κB and STAT3 reporter systems, alongside high-throughput screening of 2,820 bacterial isolates (94 per subject). Despite minimal differences in overall microbial composition, faecal water from patients with ulcerative colitis (UC) and Crohn’s disease (CD) significantly increased NF-κB activity under both basal and TNFα-stimulated conditions. Non-IBD controls harboured a higher proportion of suppressive bacterial isolates (21.52%) compared to UC (5.64%) and CD (2.45%), whereas activating isolates were enriched in IBD (UC: 34.47%; CD: 39.36% vs. non-IBD: 5.35%). Similar trends were observed for IL-23-mediated STAT3 activation.

Integration of metagenomic and pathway-level analyses identified 24 KEGG pathways associated with inflammatory signalling and faecal lipocalin-2, as well as amino acid, carbohydrate, lipid, cofactor, and nucleotide metabolism. These findings demonstrate that the IBD microbiome is functionally reprogrammed toward pro-inflammatory activity, independent of compositional changes. Defining patient-specific MIF profiles may provide a framework for precision microbiome-based therapeutic strategies in IBD.

## Main text introduction

Inflammatory Bowel Disease (IBD), encompassing Crohn’s disease (CD) and ulcerative colitis (UC), represents a chronic and relapsing inflammatory condition of the gastrointestinal tract with significant clinical morbidity and economic impacts worldwide^1^. The global incidence of IBD continues to rise, particularly in newly industrialized countries, suggesting that environmental factors play a critical role in disease pathogensis^1^ . Large-scale genome-wide association studies (GWAS) have identified over 200 genetic loci associated with IBD susceptibility identifying important molecular pathways contributing to IBD, yet these genetic factors explain only a fraction of disease risk, underscoring the importance of environmental contributions to disease pathogenesis^2^.

Environmental factors including diets high in animal fat and low in fibre, smoking, antibiotics and medication usage, are associated with increased risk of IBD^2^. Environmental risk factors associated with IBD have been postulated to affect the microbiota and thereby influence mucosal inflammation. The causal relationship between gut microbiota and IBD pathogenesis has been demonstrated in germ-free mouse models, where disease manifestation requires the presence of intestinal bacteria^3^. Furthermore, transfer of microbiota from IBD patients to germ-free mice can recapitulate aspects of intestinal inflammation, providing direct evidence for the pathogenic potential of the IBD-associated microbiome^4^.

Culture-independent metagenomic analyses have consistently revealed compositional alterations in the IBD microbiome, characterized by reduced microbial diversity, decreased abundance of obligate anaerobes (particularly *Faecalibacterium prausnitzii* and other butyrate-producing Firmicutes), and increased representation of facultative anaerobes, especially Proteobacteria such as *Escherichia coli*^3, 5, 6^. However, the functional consequences of these compositional changes remain incompletely understood. Recent metabolomic studies have identified alterations in the fecal metabolite landscape of IBD patients, including reductions in short-chain fatty acids, secondary bile acids, sphingolipids, and vitamins, alongside increases in primary bile acids, amino acids, and acylcarnitines^7, 8^. Yet, these observations are often confounded by disease-related factors such as inflammation-induced malabsorption, prior surgical interventions, and medication effects^9, 10^.

Microbial metabolites serve as potentially critical signaling molecules that mediate host-microbiota interactions and regulate intestinal immune homeostasis^11–15^. However, our understanding of how these metabolites collectively shape the immune tone of the gut what we term the microbial immunomodulatory function (MIF) remains in its infancy. Two signaling pathways, NF-κB (nuclear factor-κB) and Jak (Janus kinase)/STAT (Signal Transducer and Activator of Transcription) signalling pathways, are particularly relevant to IBD pathogenesis^11^ and represent major therapeutic targets. Within intestinal immune signalling, NF-κB functions as a central upstream regulator of pro-inflammatory responses, integrating signals from microbial products, pattern recognition receptors, cytokines, and cellular stress pathways. Activation of NF-κB downstream of Toll-like receptors (TLRs), TNF receptor signalling, and IL-1 family cytokines drives expression of key inflammatory mediators including TNF, IL-6, IL-8, and IL-23, thereby amplifying mucosal immune activation and epithelial stress responses. Within JAK/STAT signaling, STAT3 activation downstream of IL-23 signaling is particularly relevant as it promotes Th17 cell differentiation and intestinal inflammation as well as IL-17 production^16^. To achieve immune homeostasis, a degree of proinflammatory tone must be maintained to prevent infection, while tolerance to commensal microbes is critical to prevent aberrant inflammation. Current IBD therapeutics, including corticosteroids, anti-TNF agents, IL-23 inhibitors and JAK inhibitors, directly or indirectly target these pathways to control inappropriately activated immune responses.

Despite the recognized importance of the microbiome in IBD, a critical gap exists in our understanding of how microbial communities functionally modulate these key inflammatory pathways to maintain homeostasis or drive aberrant immune response. Previous studies have largely focused on compositional profiling or targeted metabolite analysis^17^, but few have directly assessed the integrated immunomodulatory capacity of the gut microbiome in IBD patients. Moreover, the relationship between microbial composition, functional metabolic output, and immunomodulatory effects on host signaling pathways remains poorly characterized.

In this study, we employed a multi-faceted approach combining shotgun metagenomics, high-throughput functional screening of bacterial isolates, and pathway-level functional analysis to characterize the microbial immunomodulatory landscape in IBD. We hypothesized that IBD patients harbor microbiota with diminished capacity to suppress inflammatory signaling and enhanced capacity to activate NF-κB and STAT3 pathways, independent of compositional differences. Furthermore, we sought to identify specific microbial species and metabolic pathways associated with these functional alterations. Our findings reveal that IBD patients’ faecal microbiota lack the ability to suppress IL-23-driven STAT3 and TNF-mediated NF-κB activation compared to healthy controls, and that this functional deficit is independent of inflammation status or overall microbial composition. These results suggest that characterizing patient-specific MIF profiles may provide greater insight into the microbial contribution to patients’ disease phenotype and enable microbiome guided precision medicine approaches for IBD treatment.

## MATERIALS AND METHODS

### Study cohort and sample collection

Stool samples were collected from 30 subjects: 10 non-IBD controls, 10 patients with ulcerative colitis (UC), and 10 patients with Crohn’s disease (CD). All samples were obtained through the Mater Inflammatory Bowel Disease Biobank in accordance with protocols approved by the Mater Health Services Human Research Ethics Committee (HREC 2016001782 and HREC/MML/7327). Written informed consent was obtained from all participants. Upon collection, stool samples were immediately transferred to an anaerobic chamber (Coy Laboratory Products) and stored in anaerobic glycerol stocks at -80°C until further processing.

### Faecal water preparation

Faecal water was prepared by homogenizing stool samples in sterile phosphate-buffered saline (PBS) at a ratio of 1 g stool to 10 ml PBS. Samples were vortexed thoroughly to ensure complete homogenization, then centrifuged at maximum speed (≥16,000 × g) for 10 minutes at 4°C. The supernatant was collected and filter-sterilized through 0.22 μm filters to remove bacteria and solid matter. Faecal water was tested at a final concentration of 10% (v/v) in cell culture assays as described below.

### DNA extraction, shotgun metagenomic sequencing, and analysis

Genomic DNA was extracted from stool samples using a repeated bead-beating protocol combined with automated column-based purification. Briefly, approximately 200 mg of stool was subjected to mechanical lysis using 0.1 mm zirconia/silica beads in a bead beater (three cycles of 60 seconds at maximum speed with 1-minute cooling on ice between cycles). Following bead beating, DNA was purified using the Maxwell RSC Blood DNA Kit (Promega) on a Maxwell RSC instrument according to the manufacturer’s instructions. DNA concentration and quality were assessed using a NanoDrop spectrophotometer and Qubit fluorometer.

Extracted DNA samples were submitted to the Australian Centre for Ecogenomics at the University of Queensland for library preparation and sequencing. Sequencing libraries were constructed using the Illumina Nextera XT DNA Library Preparation Kit following the manufacturer’s protocol. Shotgun metagenomic sequencing was performed on the Illumina NextSeq 500 platform using 150 bp paired-end reads, with a target sequencing depth of 5 Gbp per sample.

The deduplication of the metagenomic reads was carried out using BBmap/clumpify.sh (version 39.08) . Host contamination was removed using minimap2 (version 2.28) with the reference human genome hg38. The low-quality metagenomic reads were then removed with fastp (version 0.23.4). The taxonomic profiling was performed by Kraken2 (version 2.1.3) with the Genome Taxonomy Database (GTDB release 220.0 . The relative abundances of microbial features were estimated by Bracken2 (version 2.9) [6]. Microbiome functional pathways were profiled using HUMAnN (version 3.9).

Downstream diversity and statistical analyses were performed in R using the packages tidyverse, vegan, ggpubr, ggrepel, and pheatmap. Alpha diversity was assessed using Shannon diversity and species richness. Group-level differences were tested using Kruskal– Wallis tests, followed by pairwise Wilcoxon rank-sum tests with Benjamini–Hochberg correction for multiple comparisons. Beta diversity was assessed using Bray–Curtis and Jaccard distances and visualised using both unconstrained PCoA and constrained PCoA/CAP analyses. Species vectors were fitted to ordinations using envfit, with the top eight species vectors visualised to indicate taxa contributing most strongly to sample separation. The top 50 most abundant species were visualised using heatmaps of log-transformed relative abundances, with samples annotated by their respective metadata groups.

Statistical analysis was completed in R version 4.4.2. The Mann-Whitney-Wilcoxon test and the Kruskal-Wallis test were applied in the comparisons of feature abundance and alpha diversities between groups. Adonis based on Bray-Curtis distances was used to investigate and rank the effect of the metadata factors on overall microbial composition. MaAsLin3 was performed to determine the multivariable association between microbial signatures and functional pathways and clinical data. P values were adjusted for multiple testing where appropriate, using the Benjamini-Hochberg method. The plots were constructed mainly in the ggplot2 package.

### Bacterial isolation and culture

For bacterial isolation, stool samples from all 30 subjects were processed under anaerobic conditions. Each sample was serially diluted in pre-reduced PBS and plated on M10 agar medium, a formulation designed to support growth of diverse gut commensals^18^. M10 medium composition included: peptone (5 g/L), yeast extract (2.5 g/L), glucose (1 g/L), cellobiose (1 g/L), soluble starch (1 g/L), K HPO (0.45 g/L), NaCl (9 g/L), MgSO ·7H O (0.09 g/L), CaCl ·2H O (0.09 g/L), hemin (10 mg/L), vitamin K1 (1 mg/L), L-cysteine (0.5 g/L), and agar (15 g/L for solid medium). The medium was prepared anaerobically and reduced with L-cysteine immediately before use.

Plates were incubated anaerobically at 37°C for 48-72 hours. From each subject, 94 morphologically distinct colonies were randomly selected and individually inoculated into 96-well deep-well plates containing 1 ml M10 broth per well. Cultures were incubated anaerobically at 37°C for 72 hours. Following incubation, cultures were centrifuged at 4,000 × g for 10 minutes, and cell-free supernatants were collected and filter-sterilized through 0.22 μm filters. Supernatants were stored at -80°C until functional screening. In total, 2,820 bacterial isolates (94 isolates × 30 subjects) were obtained and screened.

### NF-**_κ_**B reporter cell assays

NF-κB activity was assessed using LS174T cells (ATCC) stably transfected with an NF-κB-responsive firefly luciferase reporter construct, as previously described^11, 15^. Cells were maintained in DMEM supplemented with 10% fetal bovine serum, 100 U/ml penicillin, 100 μg/ml streptomycin, and 400 μg/ml G418 for selection.

For faecal water experiments, cells were pre-treated with 10% (v/v) faecal water for 30 minutes, followed by stimulation with recombinant human TNF-α (50 ng/ml; R&D Systems) for 4 hours. For bacterial supernatant screening, cells were pre-treated with 10% (v/v) filter-sterilized culture supernatants for 30 minutes, followed by TNF-α stimulation (50 ng/ml) for 4 hours. Control wells received medium alone, medium + TNF-α, or medium + faecal water/supernatant without TNF-α.

Following stimulation, luciferase activity was measured using the Pierce Firefly Luc One-Step Glow Assay Kit (Thermo Fisher Scientific) according to the manufacturer’s instructions and luminescence was measured immediately. Results were normalized to medium + TNF-α controls (set to 100%) for each plate.

Bacterial isolates were classified as: (1) suppressive if they reduced TNF-α-induced NF-κB activation by ≥50% compared to medium + TNF-α control; (2) activating if they increased NF-κB activation >20% above the medium + TNF-α control; or (3) neutral if they had no significant effect (within ±20% of control). The proportion of isolates in each category was calculated for each subject.

### STAT3 reporter cell assays

STAT3 activation was assessed using HEK-Blue IL-23 cells (InvivoGen), which are HEK293 cells stably transfected with human IL-23 receptor, IL-12Rβ1, and a STAT3-responsive secreted embryonic alkaline phosphatase (SEAP) reporter. Cells were maintained in DMEM supplemented with 10% fetal bovine serum, 100 U/ml penicillin, 100 μg/ml streptomycin, 100 μg/ml Normocin, and selection antibiotics (Zeocin and Blasticidin) according to the manufacturer’s instructions.

For assays, cells were seeded in 96-well plates at 5 × 10 cells per well. Cells were pre-treated with 10% (v/v) faecal water or bacterial culture supernatants for 30 minutes, followed by stimulation with recombinant human IL-23 (5 ng/ml; R&D Systems) for 18 hours. Following stimulation, 20 μl of cell culture supernatant was transferred to a new plate and mixed with 180 μl of QUANTI-Blue detection reagent (InvivoGen). After 30 minutes of incubation at 37°C, SEAP activity was measured by reading optical density at 630 nm using a plate reader.

Results were normalized to medium + IL-23 controls (set to 100%) for each plate. Isolates were classified as suppressive, activating, or neutral using the same criteria as for NF-κB assays.

### Enzyme-Linked Immunosorbent Assays (ELISAs)

To determine whether inflammatory cytokines were present in faecal water, we measured IL-23, IL-6, IL-8, and TNF-α using commercially available ELISA kits. IL-23 and lipocalin-2 were measured using DuoSet ELISA kits from R&D Systems. IL-6, IL-8, and TNF-α were measured using ELISA MAX Deluxe kits from BioLegend. All assays were performed according to the manufacturers’ instructions. Briefly, 96-well plates were coated with capture antibody overnight at 4°C, blocked with reagent diluent, and incubated with standards or samples (faecal water diluted 1:2 or 1:10 in reagent diluent) for 2 hours at room temperature. After washing, detection antibody was added for 2 hours, followed by streptavidin-HRP and TMB substrate. Reactions were stopped with sulfuric acid, and absorbance was measured at 450 nm with wavelength correction at 570 nm.

### Statistical Analysis

All statistical tests were two-sided, and p-values < 0.05 (or FDR < 0.25 for metagenomic analyses) were considered statistically significant. Data visualization was performed using ggplot2, pheatmap, and other R package and Graphpad Prism

## RESULTS

### Microbial composition shows modest differences between IBD patients and controls

To establish the baseline microbial composition across samples, we performed shotgun metagenomic sequencing on stool samples from non-IBD controls (n=10), UC (n=10), and CD patients (n=10). Alpha diversity analysis revealed no significant differences in overall microbial diversity between IBD and non-IBD groups. Shannon diversity, species evenness, and observed richness were comparable across groups (Figure 1A-B, Supplementary Figure 1A-B).

**Figure 1.**
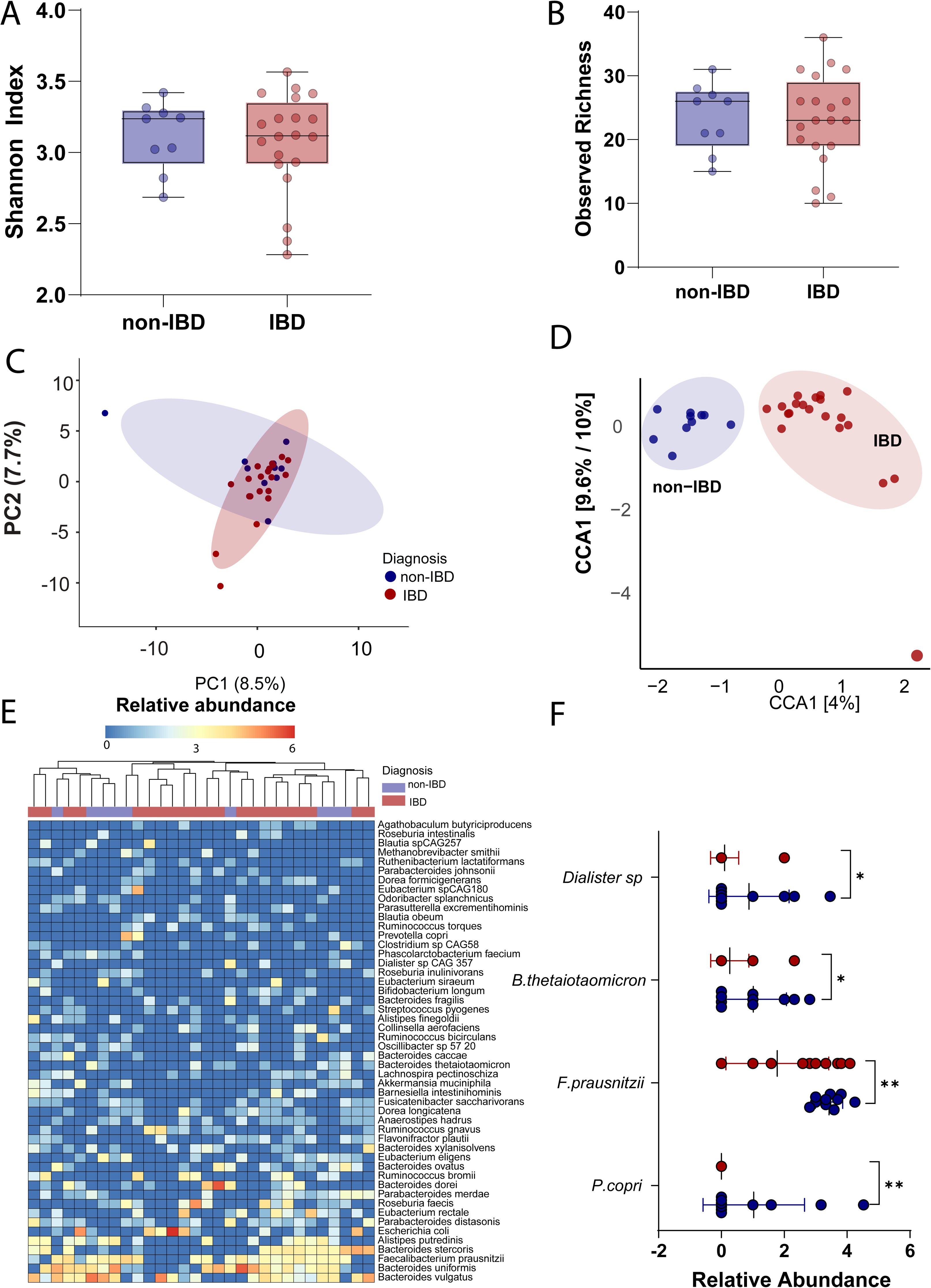
Shotgun metagenomic sequencing reveals modest compositional differences between non-IBD controls and IBD patients, with selective depletion of key commensal species. (A) Shannon Diversity Index in stool from non-IBD controls and IBD patients. (B) Observed species richness (number of detected species per sample) across non-IBD and IBD.(C) Unsupervised principal component analysis (PCA) of species-level relative abundances based on Bray-Curtis dissimilarity. Each symbol represents an individual subject, coloured by disease group . Axes indicate the percentage of total variance explained by each principal component. (D) Supervised canonical correspondence analysis (CCA) with disease status included as a constraining variable. (E) Relative abundance of selected differentially abundant species in stool from non-IBD and IBD subjects. (F) Relative abundance of *Prevotella copri*, *Bacteroides thetaiotaomicron*, *Dialister* and *Faecalibacterium prausnitzii* in stool from non-IBD (blue) and IBD (red) subjects. Each circle represents an individual subject. Statistical comparisons were performed using the Wilcoxon rank-sum (Mann– Whitney U) test. Significance is denoted as follows: * p < 0.05; ** < 0.01; *** < 0.001; **** < 0.0001; ns, not significant.

Unsupervised principal component analysis (PCA) based on Bray-Curtis dissimilarity demonstrated substantial inter-individual variation with no clear separation between groups (Figure 1C, Supplementary Figure 1C). In contrast, supervised canonical correspondence analysis (CCA) suggested modest compositional differences when disease diagnosis was included as a constraining variable (Figure 1D, Supplementary Figure 1D).

At the species level, several taxa differed significantly between groups (Figure 1E). *Faecalibacterium prausnitzii*, a key anti-inflammatory commensal, was significantly reduced in IBD patients compared to controls (p<0.05) (Figure 1F, Supplementary Figure 1E). Similarly, *Prevotella copri*, *Bacteroides thetaiotaomicron*, and *Dialister* species were enriched in non-IBD controls. This is consistent with numerous previous reports and validates our cohort as representative of typical IBD-associated dysbiosis^19, 20^. Despite these species-level differences, the overall microbial profile in our cohort showed relatively modest alterations compared to some previous reports. This may reflect several confounding factors including variation in disease activity, medication use (particularly immunosuppressants and biologics), disease behavior (inflammatory vs. stricturing vs. penetrating in CD), and disease location^21^. Importantly, faecal lipocalin-2, a marker of intestinal inflammation, was significantly elevated in both IBD subject groups compared to non-IBD controls (Supplementary Figure 2A-B), confirming the presence of active or recent inflammation in the patient cohort.

### Faecal microbiota from IBD patients exhibit enhanced NF-**_κ_**B-activating capacity independent of cytokine content

Given the modest compositional differences observed between groups, we next assessed whether the functional immunomodulatory capacity of the microbiome differed between IBD patients and non-IBD controls. To do this, cell-free faecal water from each subject was tested for its ability to modulate NF-κB activation using LS174T reporter cells.

Faecal water from both UC and CD patients significantly increased NF-κB activity under basal (unstimulated) conditions compared to non-IBD controls (p<0.05) (Figure 2A). This effect was maintained and further enhanced in the presence of TNF-α (50 ng/ml) (Figure 2B), suggesting that IBD-associated microbial metabolites can potentiate inflammatory signalling.

**Figure 2.**
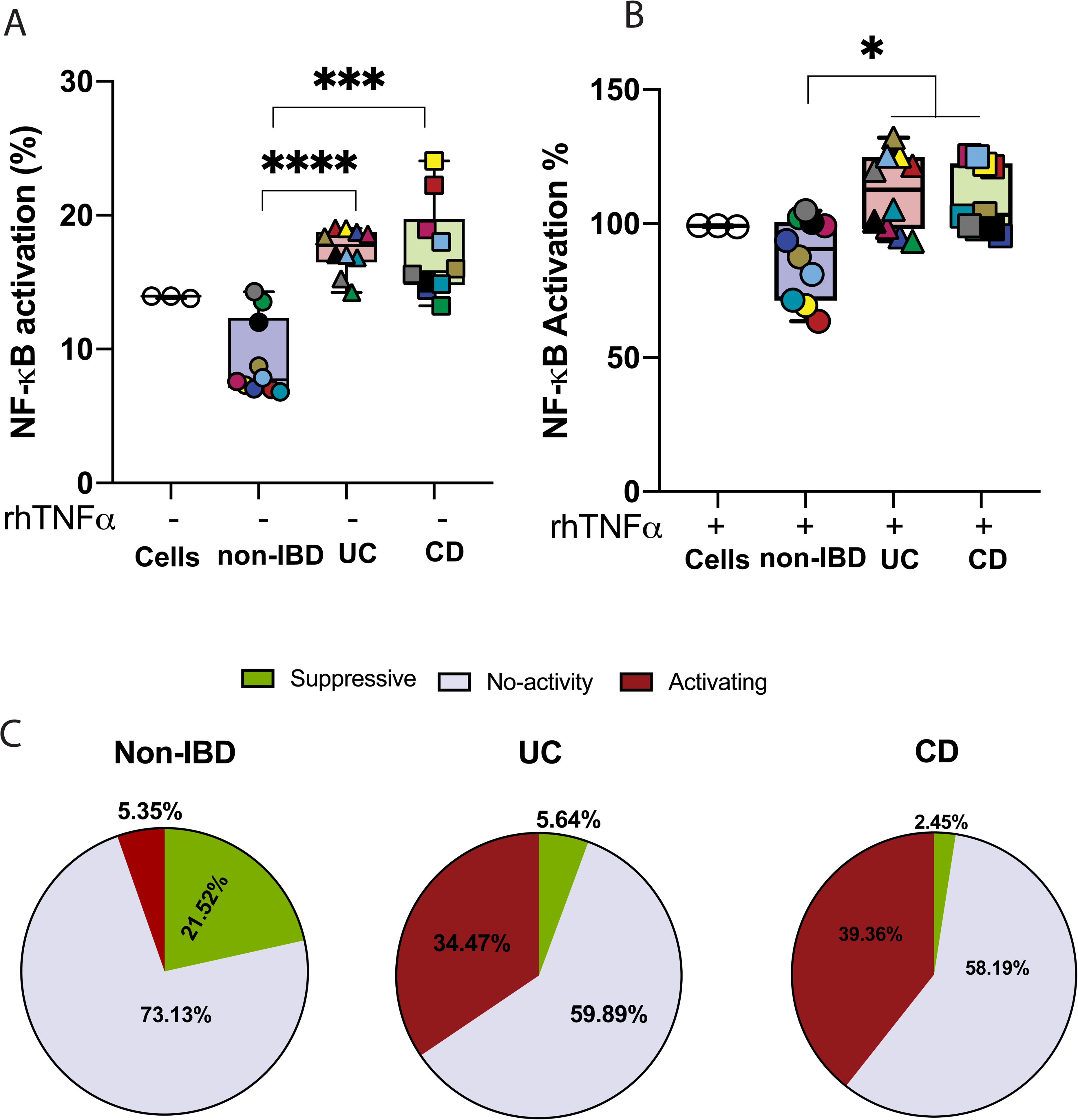
Faecal water from IBD patients activates NF-_κ_B signalling, and IBD-associated bacterial isolates show a marked shift toward pro-inflammatory immunomodulatory function. **(A)** NF-κB activation in LS174T NF-κB reporter cells treated with cell-free faecal water from non-IBD controls, UC patients, and CD patients under basal (unstimulated) conditions. NF-κB activity is expressed as % to TNF treated cells (B) NF-κB activation in LS174T reporter cells treated with faecal water in the presence of TNF-α co-stimulation (50 ng/mL). Each symbol represents an individual subject (C) Proportion of bacterial isolates classified as NF-κB-suppressive, neutral, or NF-κB-activating in non-IBD controls, UC patients, and CD patients. Suppressive isolates were defined as those reducing TNF-α-induced NF-κB activation by ≥50%; activating isolates were those enhancing NF-κB activation above the TNF-α-stimulated control. Statistical comparisons by chi-squared test. Significance is denoted as follows: *p < 0.05; ** < 0.01; *** < 0.001; **** < 0.0001; ns, not significant.

Substantial inter-individual variation was observed within each group, with some IBD samples exhibiting particularly strong NF-κB-activating capacity. To determine whether this effect was driven by residual inflammatory cytokines, levels of IL-23, IL-6, IL-8 and TNF-α were measured in faecal water. None were detectable (Supplementary Figure 3A-D), suggesting that the observed NF-κB activation was unlikely to be explained by residual host inflammatory cytokines or chemokines and is more likely attributable to microbial-derived factors.

### IBD patients harbor fewer suppressive and more activating bacterial isolates

To identify contributors of individual bacterial strains to the observed functional differences, we isolated 94 colonies per subject (2,820 total isolates) and screened their cell-free supernatants for effects on NF-κB activation. Isolates were classified as suppressive (≥50% reduction in TNF-α-induced NF-κB), activating (enhanced activation), or neutral. Non-IBD controls harbored a significantly higher proportion of suppressive isolates (21.52%) compared to UC (5.64%) and CD (2.45%) (p<0.001) (Figure 2C, Supplementary Figure 4A-C). In contrast, activating isolates were markedly enriched in IBD patients (UC: 34.47%, CD: 39.36%) relative to controls (5.35%) (p<0.001).

These findings demonstrate a pronounced shift in the functional composition of the microbiome in IBD, characterized by depletion of suppressive activity and enrichment of pro-inflammatory activity. Notably, this functional shift was substantially greater than the compositional differences observed at the species level.

### STAT3 activation mirrors NF-**_κ_**B functional alterations

To determine whether the observed functional changes extended beyond NF-κB signalling, we next assessed STAT3 activation using HEK-Blue IL-23 reporter cells. Consistent with the NF-κB findings, faecal water from IBD patients showed an increased capacity to activate STAT3 signalling in the presence of IL-23 (5 ng/ml) compared to non-IBD controls (Figure 3A-B). Analysis of bacterial isolates revealed a similar pattern. Non-IBD controls harboured a higher proportion of STAT3-suppressive isolates, whereas IBD patients were enriched for STAT3-activating isolates, although the magnitude of this difference was less pronounced than that observed for NF-κB (Figure 3C, Supplementary Figure 4D-F).

**Figure 3.**
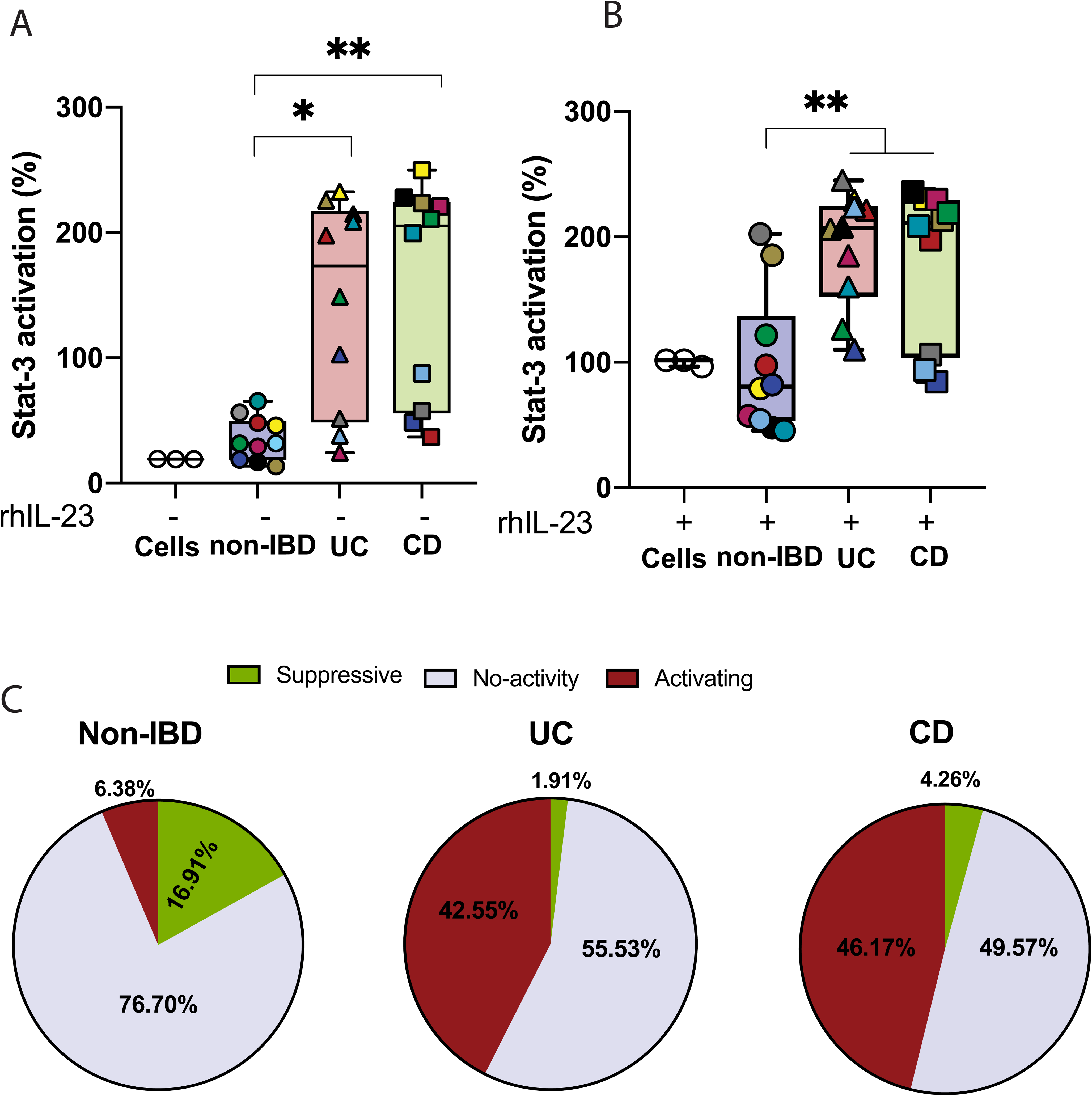
STAT3 activation by faecal water and bacterial isolates mirrors NF-_κ_B findings. (A) STAT3 activation in HEK-Blue IL-23 reporter cells treated with cell-free faecal water from non-IBD controls, UC patients, and CD patients in the absence (A) or presence (B) of IL-23 stimulation (5 ng/ml). STAT3 activity is expressed as % to IL-23 treated cells Each symbol represents an individual subject. Statistical comparisons by Kruskal-Wallis test with Dunn’s post-hoc correction. (C) Proportion of bacterial isolates classified as STAT3-suppressive, neutral, or STAT3-activating in non-IBD controls, UC patients, and CD patients, tested on HEK-Blue IL-23 reporter cells in the presence of IL-23 stimulation. Significance is denoted as follows: *p < 0.05; ** < 0.01; *** < 0.001; **** < 0.0001; ns, not significant.

Together, these findings demonstrate that the functional alterations observed in the IBD microbiome are not restricted to a single pathway but reflect a broader shift toward pro-inflammatory activity across key inflammatory signalling pathways.

### Patients stratify into distinct microbial immunomodulatory function subtypes

Integrating NF-κB and STAT3 activation profiles of faecal water from each subject revealed four distinct microbial immunomodulatory function (MIF) subtypes: (1) low activity in both pathways, (2) NF-κB-dominant activity, (3) STAT3-dominant activity, and (4) dual activation (Supplementary Figure 5A). Non-IBD controls predominantly clustered within the low-activity subtype, whereas IBD patients were distributed across the remaining subtypes, highlighting substantial functional heterogeneity.

This stratification demonstrates that functional differences in the microbiome are not fully captured by clinical diagnosis alone. While some patients exhibited selective NF-κB or STAT3 activation, others showed activation across both pathways, indicating distinct underlying inflammatory profiles. These findings suggest that microbial immunomodulatory function may provide a framework for patient stratification and could inform pathway-targeted therapeutic strategies.

### Integration of metagenomics with microbial immunomodulatory function identifies species associated with inflammatory activity

To identify the specific microbial drivers of the observed immunomodulatory phenotypes, we performed MaAsLin3 associating species abundance with low inflammatory pathway activity. Notably, *Anaerostipes sp905215045* , *Roseburia sp019411365* and *Enterocloster sp900543885* exhibited the strongest associations with the NF-κB suppressive profile while *Akkermansia muciniphila* was positively associate with both NF-κB and STAT3 suppression. These taxa were less prevalent in samples with high NF-κB or STAT3 activity, suggesting that depletion of these commensal bacteria may contribute to the pro-inflammatory faecal water profiles observed in UC and CD patients (Figure 4).

**Figure 4.**
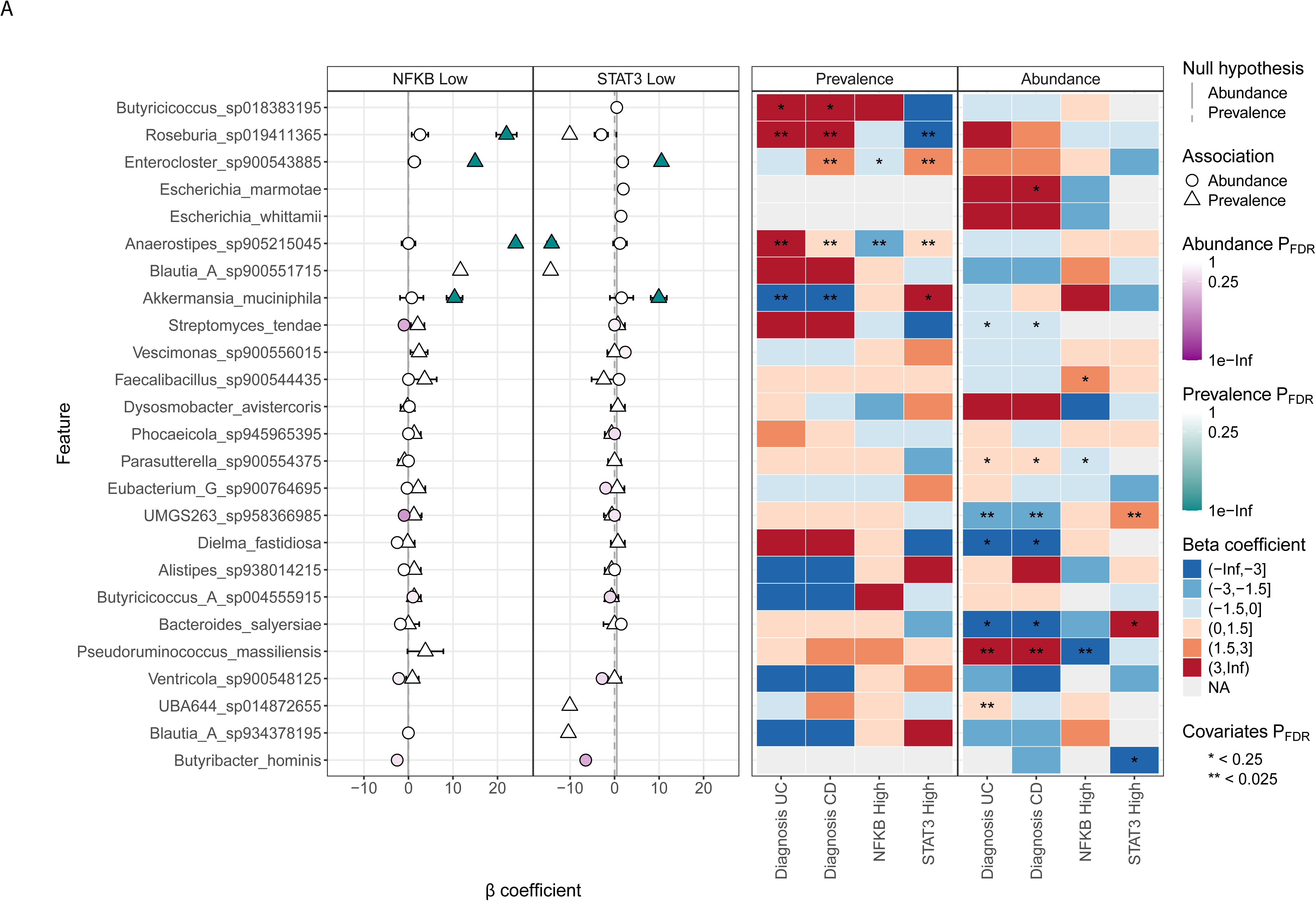
Integration of metagenomic data with microbial immunomodulatory function reveals species-level associations. (A) Dot plots showing associations between microbial species and low NF-κB activity (left panel) or low STAT3 activity (right panel). Green triangles indicate significant prevalence associations (species presence/absence); open circles indicate abundance associations (species relative abundance). Error bars represent 95% confidence intervals. (B) Heatmaps showing species associations with UC diagnosis, CD diagnosis, high NF-κB activity, and high STAT3 activity. Left panel shows prevalence-based associations; right panel shows abundance-based associations. Cell colors represent beta coefficients from MaAsLin2 models (blue = negative association, red = positive association). Asterisks indicate statistical significance: * FDR < 0.25, ** FDR < 0.025.

Our association analysis provides a taxonomic rationale for the functional ’MIF subtypes’ we identified. The strong association of *Akkermansia muciniphila* and *Roseburia* species with low NF-κB activity aligns with previous reports of their anti-inflammatory properties and ability to maintain gut barrier integrity^22–24^. Importantly, the significant associations found in our ’STAT3 High’ and ’NFKB High’ cohorts suggest that different bacterial communities may trigger distinct inflammatory pathways. These relationships were observed despite relatively modest differences in overall microbial composition, highlighting those specific taxa, rather than global diversity, are key determinants of immunomodulatory function. These findings support the concept of precision medicine in IBD, where microbial immunomodulatory signatures may help identify biologically distinct patient subgroups with differential inflammatory pathway activity.

### Microbial metabolic pathways are associated with inflammatory activity and immunomodulation

Multivariable association analysis identified a coherent and internally consistent set of pathway-level shifts linking inflammatory bowel disease status with host inflammatory signalling. Notably, most associations were detected in the prevalence model rather than relative abundance, indicating differences in pathway presence across individuals rather than uniform expansion within the microbiome. Both CD and UC showed strong positive associations with a cluster of de novo purine and deoxyribonucleotide biosynthesis pathways, including PWY-7229, PWY-6126, PWY-7228, PWY-6125, PWY-7222, and PWY-7220. These pathways were predominantly attributed to *Bacteroides dorei*, suggesting an increased likelihood of colonisation by strains with enhanced nucleotide biosynthetic capacity in disease. In contrast, disease status was negatively associated with PWY-5659 (GDP-mannose biosynthesis) and GLYCOGENSYNTH-PWY (glycogen biosynthesis), indicating reduced representation of carbohydrate storage and glycan precursor pathways in IBD^25^. Together, these findings suggest a shift toward growth-associated nucleotide synthesis at the expense of storage and structural metabolism^26^, consistent with adaptation to an inflamed and nutrient-variable gut environment (Figure 5A-B).

**Figure 5.**
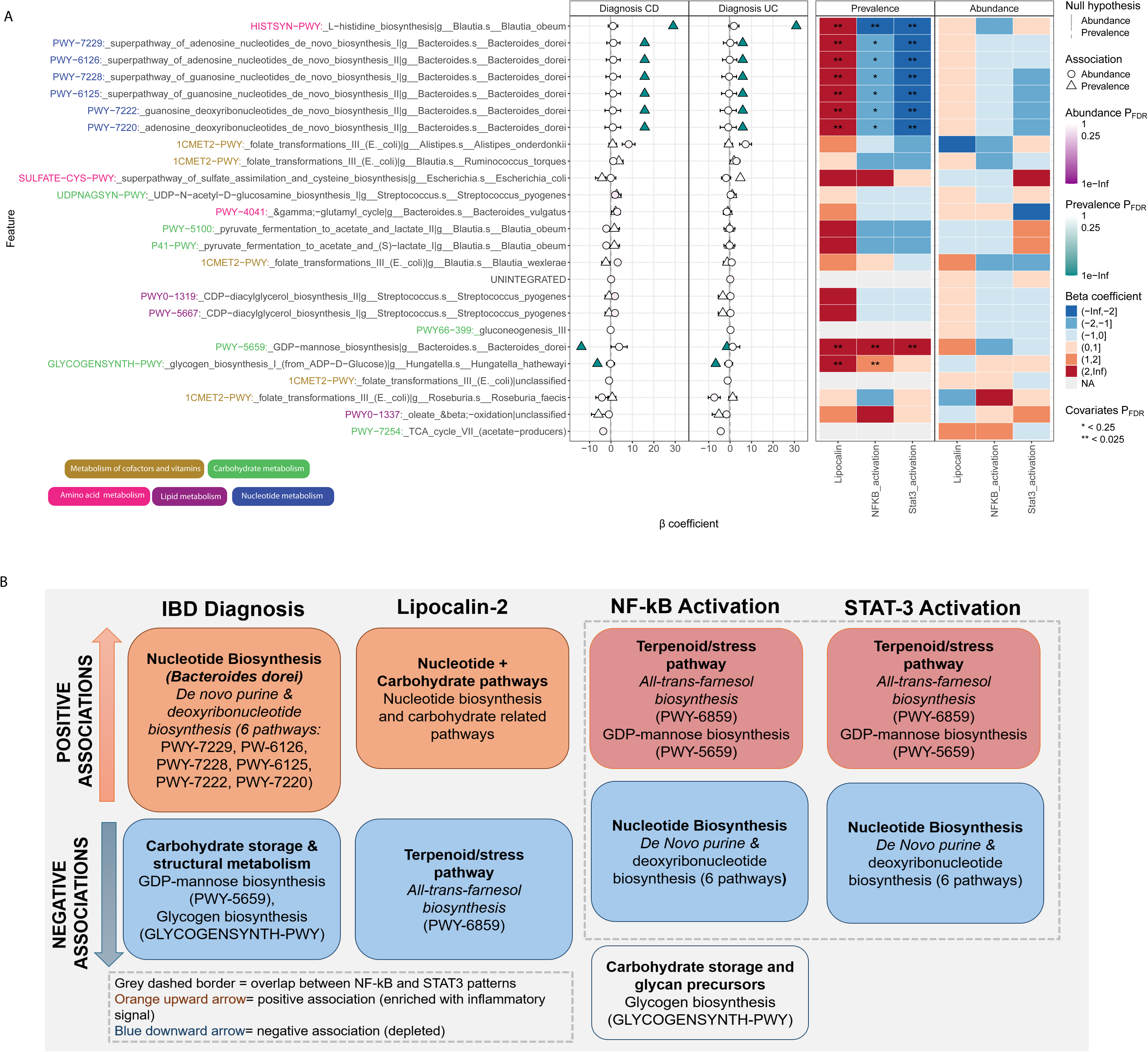
Functional metabolic pathway analysis reveals microbial metabolic pathways associated with IBD diagnosis and inflammatory pathway activation. (A) MaAsLin3 outputs showing beta coefficients for associations between KEGG metabolic pathways and clinical/functional variables (CD diagnosis, UC diagnosis, faecal lipocalin-2, NF-κB activation, STAT3 activation). Pathways are labeled with their MetaCyc IDs and contributing bacterial genera. Pathways are colored by metabolic category: amino acid metabolism, carbohydrate metabolism, lipid metabolism, metabolism of cofactors and vitamins, and nucleotide metabolism. B) Graphical summary of the major pathway-level associations identified in the prevalence model. Cell colors represent beta coefficients (blue = negative association, red = positive association). Asterisks indicate statistical significance: * FDR < 0.25, ** FDR < 0.025.

Host inflammatory signalling further refined these disease-associated patterns. Although UC and CD diagnosis were positively associated with nucleotide biosynthesis pathways, higher NF-κB activation was associated with reduced representation of these same pathways and increased representation of PWY-6859, PWY-5659 and GLYCOGENSYNTH-PWY. This divergence suggests that diagnosis and inflammatory activity are not interchangeable and likely reflect different dimensions of microbial functional variation within the cohort. This inverse relationship suggests that heightened innate immune activation is associated with reduced prevalence of metabolically intensive nucleotide synthesis pathways, alongside relative enrichment of pathways linked to stress responses and carbohydrate metabolism. STAT3 activation showed a similar but more restricted pattern, with positive associations for PWY-6859 and PWY-5659, and negative associations with the same nucleotide biosynthesis pathways. This overlap suggests coordinated selective pressures imposed by NF-κB and STAT3 signalling on microbial metabolic capacity.

In contrast, lipocalin displayed a distinct signature, with positive associations across both nucleotide biosynthesis and carbohydrate-related pathways, and a negative association with PWY-6859. Given its role in iron sequestration^27^, this pattern may reflect compensatory microbial strategies to maintain growth and storage functions under metal limitation.

Collectively, these findings demonstrate that disease status and host inflammatory pathway activity capture distinct dimensions of microbial functional variation, with several metabolic pathways exhibiting opposing associations with diagnosis and inflammatory signalling.

## DISCUSSION

This study demonstrated that gut microbiome in IBD exhibits an altered immunomodulatory function independent of compositional changes. While conventional studies have focused on characterising microbial taxonomic composition, our data show that the metagenome and its functional outputs, specifically the metabolites produced and the inflammatory pathways they modulate diverges in disease settings. We showed that non-IBD controls harboured more bacterial isolates that, following their *in vitro* cultivation with a habitat-simulating medium, produce NF-κB and STAT3 suppressive activity and with substantially fewer activating isolates, compared to IBD patients. This was consistent across both UC and CD despite their distinct anatomical distributions, characteristic patterns of inflammation and immunological drivers. This magnitude of functional divergence, in the context of relatively modest compositional change, supports a model in which dysbiosis reflects a functional collapse rather than a purely structural shift.

This apparent dissociation between microbial taxonomy and functionality is mechanistically plausible, particularly at the species-and/or strain-level of variation. For instance, different clades within the genus *Enterococcus* can confer opposing immunomodulatory properties in the context of IBD^28^. Our pathway-level analysis presented here, provides a partial resolution to this challenge by quantifying functional capabilities directly, and further reveals the metabolic redundancy and potential stability in microbial outputs during fluctuations in environmental factors such as oxygen tension, substrate availability^29^, and inflammation that further modulate microbial activity^30^.

Here, a species-level lineage within the genus *Roseburia* and *Akkermansia muciniphila* emerged as the bacterial taxa most consistently associated with suppressed inflammatory signalling, based on their significant and positive prevalence associations with the microbiota producing faecal waters most suppressive of both NF-κB and low STAT3 activity. *Enterocloster* sp. and *Anaerostipes* sp. showed strong protective prevalence associations specifically with low NF-κB activity, though *Anaerostipes* showed a negative abundance association in the STAT3 Low panel, indicating a more pathway-specific rather than broadly anti-inflammatory relationship. These findings are biologically coherent: *Roseburia* and *Enterocloster* are well-characterised butyrate producers within the Lachnospiraceae family whose fermentation end-products suppress NF-κB signalling through histone deacetylase inhibition and GPR109A-mediated activation of regulatory immune cascades^31^; *Akkermansia muciniphila* reinforces epithelial barrier integrity through mucin catabolism and the release of bioactive outer membrane proteins^32^; and *Anaerostipes* contributes to butyrate production via cross-feeding on acetate^33^. Their depletion in association with NF-κB activation is consistent with the known consequences of mucosal inflammation, which increases epithelial oxygenation and nitrate availability^34^, creating conditions that disfavour strict anaerobes while promoting expansion of inflammation-tolerant taxa. This suggests a possible self-reinforcing cycle in which inflammation drives loss of protective microbial functions, further amplifying inflammatory signalling.

The stratification of subjects into four distinct MIF subtypes: low activity in both pathways, selective NF-κB activation, selective STAT3 activation, and high activity in both reveals a dimension of IBD heterogeneity that is not detected with compositional or even clinical phenotyping. Non-IBD controls clustered predominantly in the low-activity subtype, while IBD patients distributed across all three pro-inflammatory subtypes with substantial inter-individual variation. This heterogeneity has potentially important therapeutic implications. Current IBD biologics are mechanistically selective: anti-TNF agents primarily suppress NF-κB-driven innate inflammation, IL-12/23 inhibitors target the STAT3-activating Th17 axis^35^, and JAK inhibitors provide broader inhibition across multiple STAT pathways. The existence of patients with selectively elevated NF-κB but not STAT3 or vice versa suggests that MIF profiling could serve as a functional biomarker for treatment selection, directing NF-κB-dominant patients toward anti-TNF therapy and STAT3-dominant patients toward IL-23 or JAK inhibition. Patients with activation of both pathways may represent the most refractory subgroup, potentially requiring combination biological therapy or novel agents targeting shared upstream nodes. Prospective validation of this precision medicine framework in treatment-naïve cohorts with longitudinal follow-up will be an essential next step.

Several limitations of the current study must be acknowledged. The cohort was modest in size, and the absence of stratification by disease activity, medication use, and disease behaviour introduces confounding that larger studies with detailed clinical phenotyping will need to address. The study is focused on luminal (fecal) samples rather than the mucosa-associated microbiome, which may exert more proximal effects on epithelial immune signaling . We also recognize there can be species-and/or strain-level differences in MIF^11, 36^ and the biobank of bacterial isolates used here has not yet been subjected to the level of DNA sequencing necessary to reveal any genetic bases for the variations. Despite these constraints, the convergence of evidence across three independent analytical platforms: functional isolate screening, species-level modelling^37^, and KEGG pathway analysis substantially strengthens confidence in the principal findings.

Taken together, our results support a model in which IBD is characterized by the alteration of microbial immunomodulation and offers an augmented assessment of alterations to the commensal bacteria inherent to health and IBD, beyond taxonomic criteria. Such knowledge is key if the restoration of the functional balance through targeted microbial or metabolic interventions are to be achievable with a greater level of efficacy and also repesents a promising direction for the identification of future therapeutic targets. Functional microbiome profiling, as applied here, provides a framework for linking microbial ecology to host immune responses and for translating these insights into clinically actionable strategies.

## Acknowledgement

We sincerely thank all patients and healthy volunteers who generously contributed samples for this study. We acknowledge the support of the Translational Research Institute (TRI) core facilities.

## Supplementary Figure Legends

**Supplementary Figure 1.**
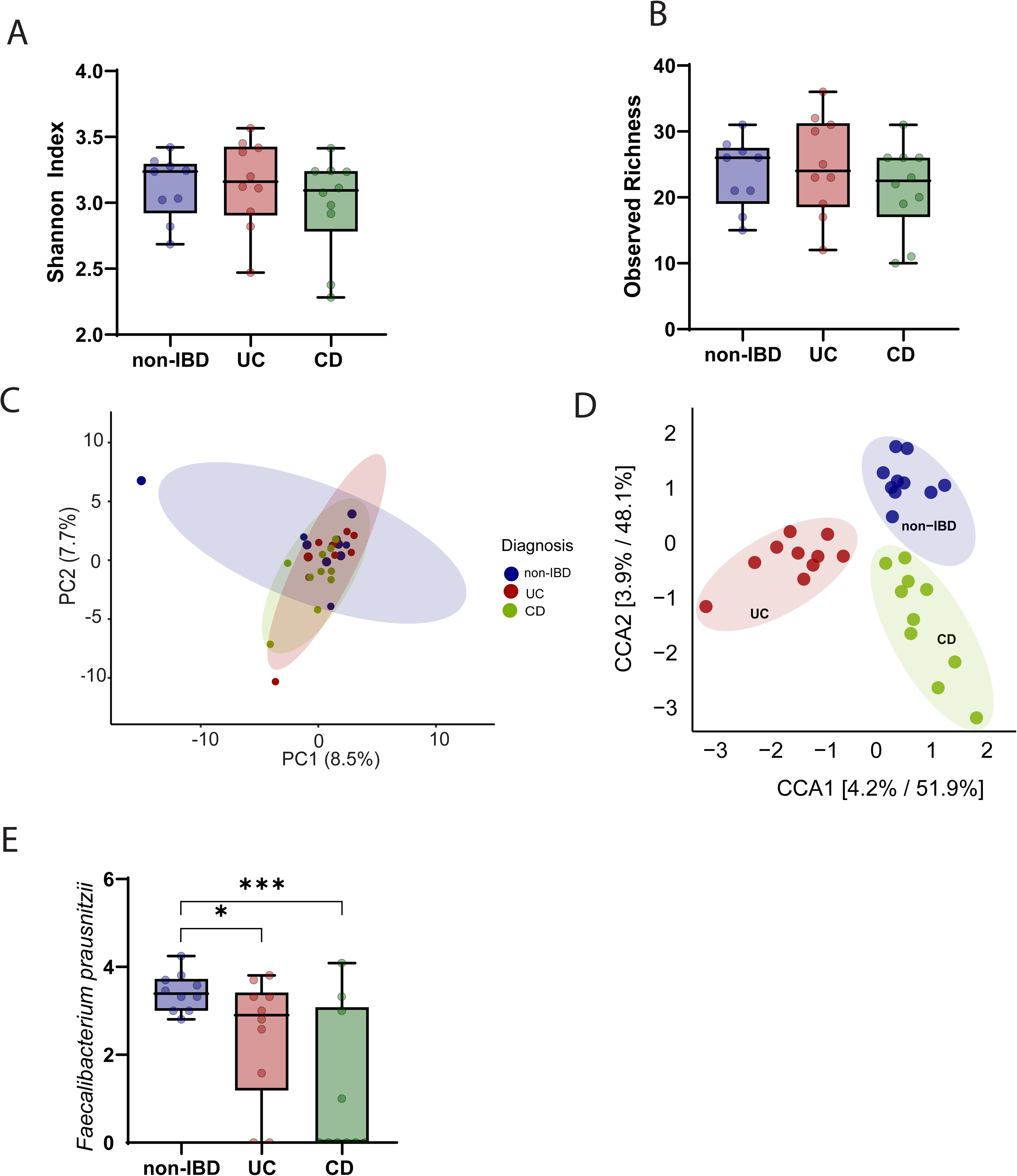
Microbial composition does not differ significantly between non-IBD, UC, and CD patients. (A) Shannon Diversity Index in stool from non-IBD, UC, and CD subjects. Each circle represents an individual subject. Statistical comparisons were performed using Kruskal-Wallis test with Dunn’s post-hoc correction; no significant differences were observed between groups. (B) Observed species richness in non-IBD, UC, and CD subjects. Each circle represents an individual subject; no significant differences were detected between groups. (C) Unsupervised principal component analysis (PCA) based on Bray-Curtis dissimilarity of species-level relative abundances from shotgun metagenomic sequencing. Each symbol represents an individual subject, coloured by group (non-IBD, UC, CD). No significant clustering by disease status was observed. (D) Supervised canonical correspondence analysis (CCA) with disease status as the constraining variable, showing subtle group-level compositional differences. (E) Relative abundance of *Faecalibacterium prausnitzii* in stool from non-IBD, UC, and CD subjects individually. *F. prausnitzii* was significantly reduced in both UC and CD patients compared to non-IBD controls. Each circle represents an individual subject. p < 0.05.

**Supplementary Figure 2.**
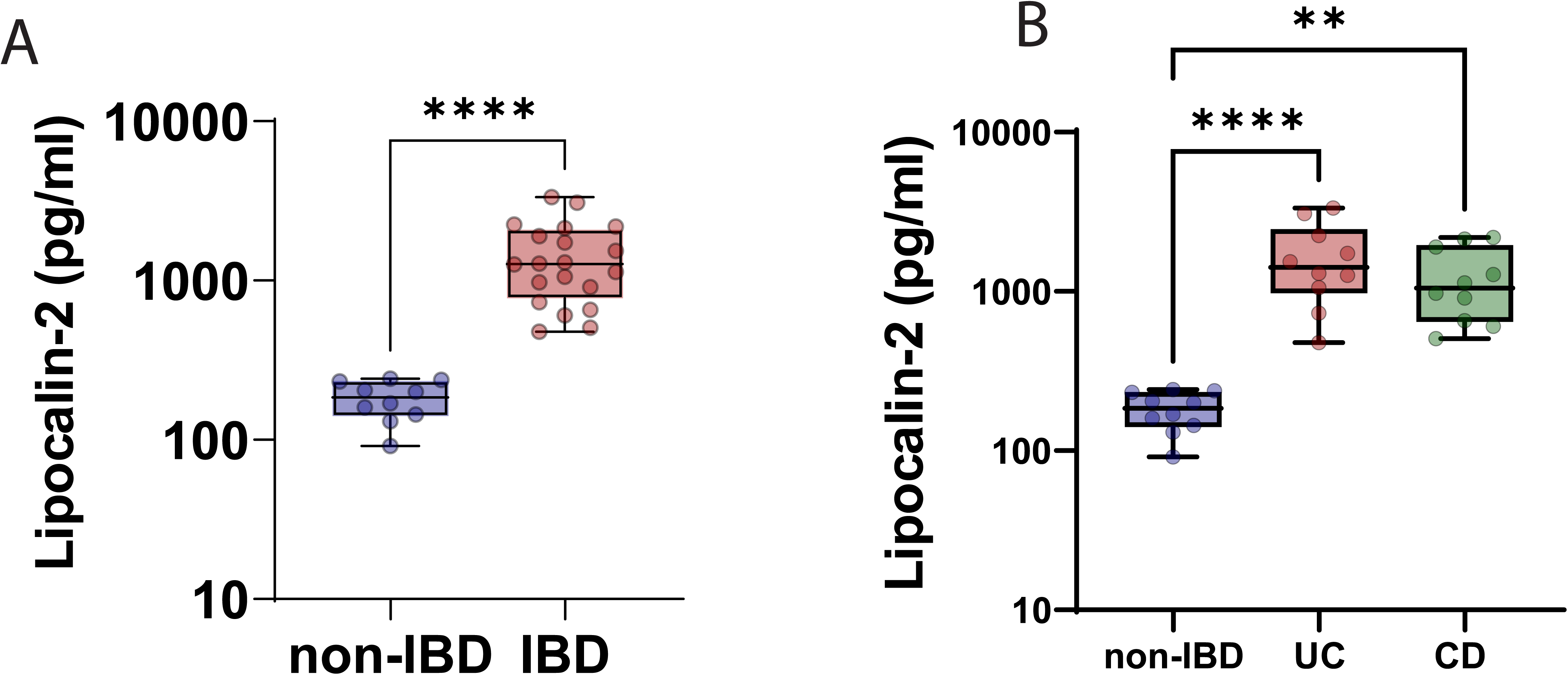
Faecal lipocalin-2 is elevated in IBD patients compared to non-IBD controls.. (A) Lipocalin-2 concentrations measured by ELISA in faecal water from non-IBD and IBD subjects. UC and CD patients are grouped together as IBD. Each circle represents an individual subject. IBD patients showed significantly higher lipocalin-2 levels compared to non-IBD controls, confirming the presence of intestinal inflammation. Statistical comparison was performed using Mann-Whitney U test. p < 0.01. (B) Lipocalin-2 concentrations stratified by IBD subtype (UC and CD). Each circle represents an individual subject. Both UC and CD patients showed elevated lipocalin-2 relative to non-IBD controls, with no significant difference between UC and CD groups.

**Supplementary Figure 3.**
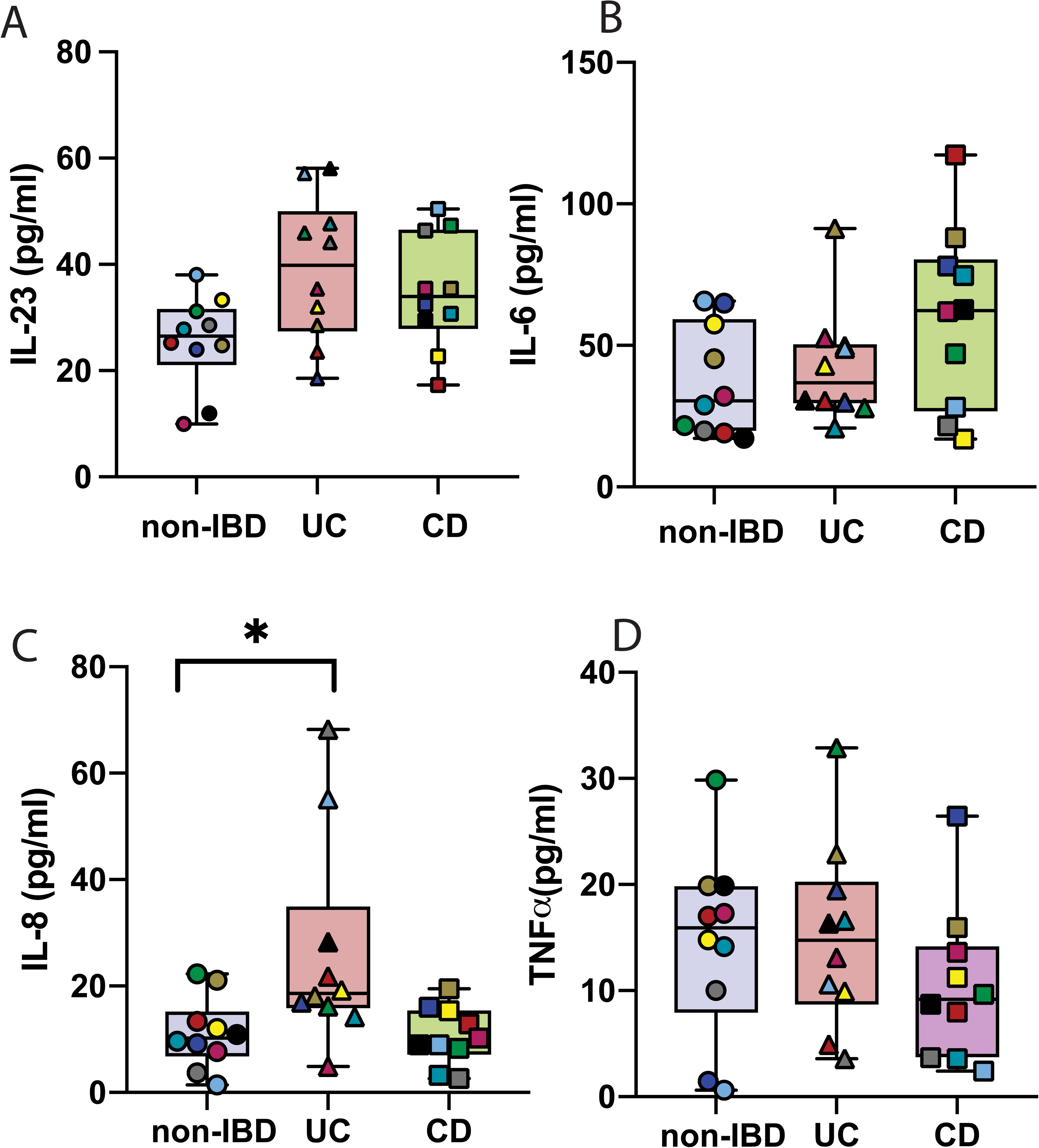
Inflammatory cytokines and chemokines are not detectable in faecal water. (A) IL-23 concentrations in faecal water from non-IBD, UC, and CD subjects measured by ELISA. Each dot represents an individual subject. IL-23 was below the limit of detection in all samples, indicating that the NF-κB and STAT3 activation observed in reporter assays was not attributable to residual cytokine content in faecal water. (B) IL-6 concentrations in faecal water from non-IBD, UC, and CD subjects. Each dot represents an individual subject. IL-6 was below the limit of detection in all samples. (C) IL-8 concentrations in faecal water from non-IBD, UC, and CD subjects. Each dot represents an individual subject. IL-8 was below the limit of detection in all samples. (D) TNF concentrations in faecal water from non-IBD, UC, and CD subjects. Each dot represents an individual subject. TNF was below the limit of detection in all samples. Dotted lines indicate the lower limit of detection for each assay.

**Supplementary Figure 4.**
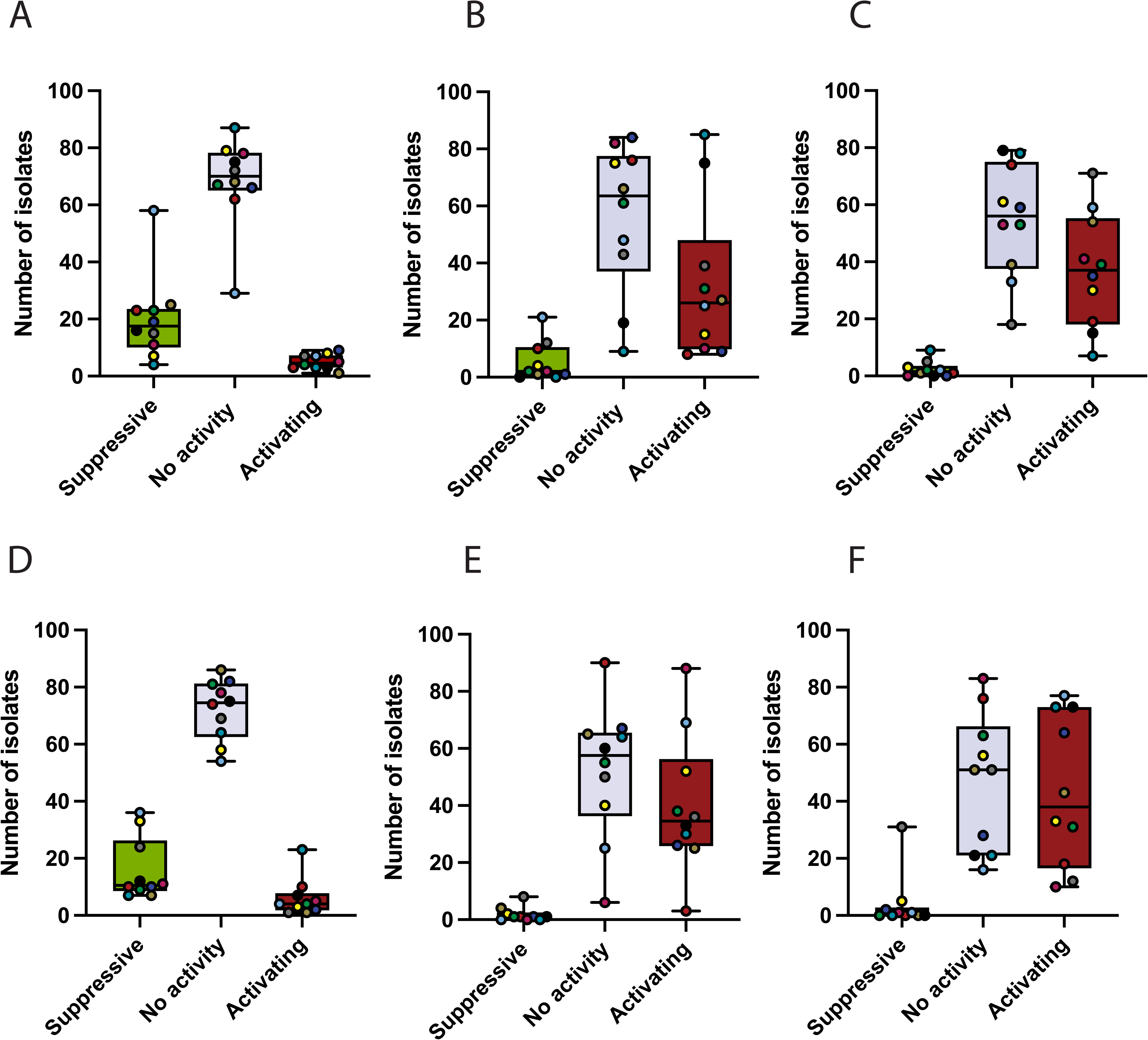
IBD patients harbour a higher proportion of NF-_κ_B-and STAT3-activating stool isolates than non-IBD subjects. (A–C) Individual-level breakdown of NF-κB immunomodulatory activity for bacterial isolates from each subject. Isolates are classified as suppressive (≥50% reduction in TNF-α-induced NF-κB activation), no activity (comparable to medium + TNF-α control), or activating (enhanced NF-κB activation above TNF-α-stimulated control). (A) Non-IBD subjects. (B) UC patients. (C) CD patients. Each coloured dot represents an individual subject, with the proportion of each functional category shown per subject. Non-IBD controls harboured significantly more suppressive isolates (21.52%) compared to UC (5.64%) and CD (2.45%), while UC and CD patients harboured significantly more activating isolates (34.47% and 39.36%, respectively) compared to non-IBD controls (5.35%). (D–F) Individual-level breakdown of STAT3 immunomodulatory activity for bacterial isolates tested on HEK-Blue IL-23 reporter cells. Isolates are classified as suppressive, no activity, or activating based on their effect on IL-23-mediated STAT3 activation. (D) Non-IBD subjects. (E) UC patients. (F) CD patients. Each coloured dot represents an individual subject. A similar trend to NF-κB was observed, with non-IBD controls harbouring more STAT3-suppressive isolates and IBD patients harbouring more STAT3-activating isolates.

**Supplementary Figure 5.**
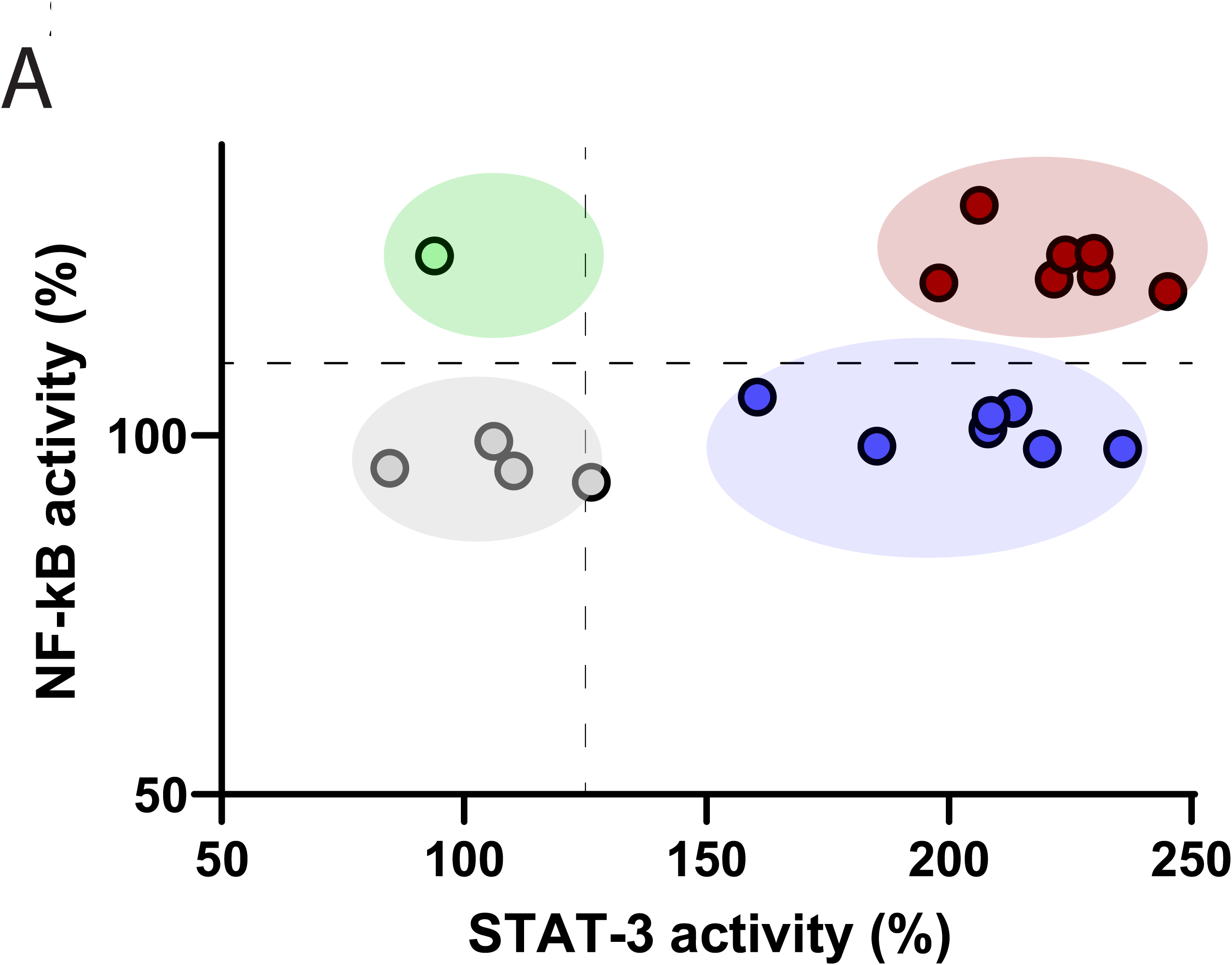
Stratification of subjects into four microbial immunomodulatory function subtypes. (A) Scatter plot showing faecal water activity on both NF-κB (x-axis) and STAT3 (y-axis) reporter cell lines for all subjects. Each circle represents an individual subject, coloured by MIF subtype: grey, low activity in both pathways; green, high NF-κB activity with low STAT3 activity; blue, high STAT3 activity with low NF-κB activity; red, high activity in both NF-κB and STAT3 pathways. Non-IBD controls predominantly cluster in the low-activity (grey) subtype, while IBD patients (UC and CD) are distributed across the three pro-inflammatory subtypes, highlighting the inter-individual heterogeneity in MIF profiles among IBD patients.

## References

1. Collaborators GBDIBD. The global, regional, and national burden of inflammatory bowel disease in 195 countries and territories, 1990-2017: a systematic analysis for the Global Burden of Disease Study 2017. Lancet Gastroenterol Hepatol 2020;5:17–30.

2. Castro F, de Souza HSP. Dietary Composition and Effects in Inflammatory Bowel Disease. Nutrients 2019;11.

3. Ni J, Wu GD, Albenberg L, et al. Gut microbiota and IBD: causation or correlation? Nat Rev Gastroenterol Hepatol 2017;14:573–584.

4. Britton GJ, Contijoch EJ, Mogno I, et al. Microbiotas from Humans with Inflammatory Bowel Disease Alter the Balance of Gut Th17 and RORgammat(+) Regulatory T Cells and Exacerbate Colitis in Mice. Immunity 2019;50:212–224 e4.

5. Zheng J, Sun Q, Zhang M, et al. Noninvasive, microbiome-based diagnosis of inflammatory bowel disease. Nat Med 2024;30:3555–3567.

6. Peter R, Zhang F, Scott G, et al. The Gut Microbiome at the Onset of Inflammatory Bowel Disease: A Systematic Review and Unified Bioinformatic Synthesis. Gastroenterology 2026;170:539–556.

7. Franzosa EA, Sirota-Madi A, Avila-Pacheco J, et al. Gut microbiome structure and metabolic activity in inflammatory bowel disease. Nat Microbiol 2019;4:293–305.

8. Heinken A, Ravcheev DA, Baldini F, et al. Systematic assessment of secondary bile acid metabolism in gut microbes reveals distinct metabolic capabilities in inflammatory bowel disease. Microbiome 2019;7:75.

9. Le Gall G, Noor SO, Ridgway K, et al. Metabolomics of fecal extracts detects altered metabolic activity of gut microbiota in ulcerative colitis and irritable bowel syndrome. J Proteome Res 2011;10:4208–18.

10. Bjerrum JT, Wang Y, Hao F, et al. Metabonomics of human fecal extracts characterize ulcerative colitis, Crohn’s disease and healthy individuals. Metabolomics 2015;11:122–133.

11. Giri R, Hoedt EC, Khushi S, et al. Secreted NF-kappaB suppressive microbial metabolites modulate gut inflammation. Cell Rep 2022;39:110646.

12. P ÓC, de Wouters T, Giri R, et al. The gut bacterium and pathobiont Bacteroides vulgatus activates NF-κB in a human gut epithelial cell line in a strain and growth phase dependent manner. Anaerobe 2017;47:209–217.

13. P ÓC, Giri R, Hoedt EC, et al. Enterococcus faecalis AHG0090 is a Genetically Tractable Bacterium and Produces a Secreted Peptidic Bioactive that Suppresses Nuclear Factor Kappa B Activation in Human Gut Epithelial Cells. Front Immunol 2018;9:790.

14. Shanahan ER, Shah A, Koloski N, et al. Influence of cigarette smoking on the human duodenal mucosa-associated microbiota. Microbiome 2018;6:150.

15. P OC, de Wouters T, Giri R, et al. The gut bacterium and pathobiont Bacteroides vulgatus activates NF-kappaB in a human gut epithelial cell line in a strain and growth phase dependent manner. Anaerobe 2017;47:209–217.

16. Jia J, Liu Y, Wang D, et al. Ulcerative colitis: signaling pathways, therapeutic targets and interventional strategies. Signal Transduct Target Ther 2026;11:51.

17. Alam MT, Amos GCA, Murphy ARJ, et al. Microbial imbalance in inflammatory bowel disease patients at different taxonomic levels. Gut Pathog 2020;12:1.

18. Eller C, Crabill MR, Bryant MP. Anaerobic roll tube media for nonselective enumeration and isolation of bacteria in human feces. Appl Microbiol 1971;22:522–9.

19. Cao Y, Shen J, Ran ZH. Association between Faecalibacterium prausnitzii Reduction and Inflammatory Bowel Disease: A Meta-Analysis and Systematic Review of the Literature. Gastroenterol Res Pract 2014;2014:872725.

20. Sokol H, Pigneur B, Watterlot L, et al. Faecalibacterium prausnitzii is an anti-inflammatory commensal bacterium identified by gut microbiota analysis of Crohn disease patients. Proc Natl Acad Sci U S A 2008;105:16731–6.

21. Galazzo G, Tedjo DI, Wintjens DSJ, et al. Faecal Microbiota Dynamics and their Relation to Disease Course in Crohn’s Disease. J Crohns Colitis 2019;13:1273–1282.

22. Martin-Gallausiaux C, Garcia-Weber D, Lashermes A, et al. Akkermansia muciniphila upregulates genes involved in maintaining the intestinal barrier function via ADP-heptose-dependent activation of the ALPK1/TIFA pathway. Gut Microbes 2022;14:2110639.

23. Daniel N, Gewirtz AT, Chassaing B. Akkermansia muciniphila counteracts the deleterious effects of dietary emulsifiers on microbiota and host metabolism. Gut 2023;72:906–917.

24. Kang X, Liu C, Ding Y, et al. Roseburia intestinalis generated butyrate boosts anti-PD-1 efficacy in colorectal cancer by activating cytotoxic CD8(+) T cells. Gut 2023;72:2112–2122.

25. Xu X, Ocansey DKW, Hang S, et al. The gut metagenomics and metabolomics signature in patients with inflammatory bowel disease. Gut Pathog 2022;14:26.

26. Kudelka MR, Stowell SR, Cummings RD, et al. Intestinal epithelial glycosylation in homeostasis and gut microbiota interactions in IBD. Nat Rev Gastroenterol Hepatol 2020;17:597–617.

27. Toyonaga T, Matsuura M, Mori K, et al. Lipocalin 2 prevents intestinal inflammation by enhancing phagocytic bacterial clearance in macrophages. Sci Rep 2016;6:35014.

28. D’Adamo GL, Chonwerawong M, Gearing LJ, et al. Bacterial clade-specific analysis identifies distinct epithelial responses in inflammatory bowel disease. Cell Rep Med 2023;4:101124.

29. Speckmann B, Ehring E, Hu J, et al. Exploring substrate-microbe interactions: a metabiotic approach toward developing targeted synbiotic compositions. Gut Microbes 2024;16:2305716.

30. Shang T, Zhang R, Liu Y, et al. Intestinal oxygen and microbiota crosstalk: implications for pathogenesis of gastrointestinal diseases and emerging therapeutic strategies. Gut Pathog 2025;17:100.

31. Bilotta AJ, Cong Y. Gut microbiota metabolite regulation of host defenses at mucosal surfaces: implication in precision medicine. Precis Clin Med 2019;2:110–119.

32. Xu W, Li A, Jing H, et al. The role of Akkermansia muciniphila in the regulation of inflammatory bowel disease: intestinal immunity and metabolism. Front Immunol 2025;16:1653472.

33. Liu D, Xie LS, Lian S, et al. Anaerostipes hadrus, a butyrate-producing bacterium capable of metabolizing 5-fluorouracil. mSphere 2024;9:e0081623.

34. Schwarzler J, Mayr L, Grabherr F, et al. Epithelial metabolism as a rheostat for intestinal inflammation and malignancy. Trends Cell Biol 2024;34:913–927.

35. Puca P, Parello S, Colantuono S, et al. Interleukin-23 Inhibitors in Inflammatory Bowel Disease. BioDrugs 2026.

36. P OC, Giri R, Hoedt EC, et al. Enterococcus faecalis AHG0090 is a Genetically Tractable Bacterium and Produces a Secreted Peptidic Bioactive that Suppresses Nuclear Factor Kappa B Activation in Human Gut Epithelial Cells. Front Immunol 2018;9:790.

37. Sudhakar P, Andrighetti T, Verstockt S, et al. Integrated analysis of microbe-host interactions in Crohn’s disease reveals potential mechanisms of microbial proteins on host gene expression. iScience 2022;25:103963.

